# Selection signatures underlying dramatic male inflorescence transformation during modern hybrid maize breeding

**DOI:** 10.1101/284109

**Authors:** Joseph L. Gage, Michael R. White, Jode W. Edwards, Shawn Kaeppler, Natalia de Leon

## Abstract

Inflorescence capacity plays a crucial role in reproductive fitness in plants, and in production of hybrid crops. Maize is a monoecious species bearing separate male and female flowers (tassel and ear, respectively). The switch from open-pollinated populations of maize to hybrid-based breeding schemes in the early 20^th^ century was accompanied by a dramatic reduction in tassel size, and the trend has continued with modern breeding over the recent decades. The goal of this study was to identify selection signatures in genes that may underlie this dramatic transformation. Using a population of 942 diverse inbred maize accessions and a nested association mapping population comprised of three 200-line biparental populations, we measured 15 tassel morphological characteristics by manual and image-based methods. Genome-wide association studies identified 242 single nucleotide polymorphisms significantly associated with measured traits. We compared 41 unselected lines from the Iowa Stiff Stalk Synthetic (BSSS) population to 21 highly selected lines developed by modern commercial breeding programs and show that tassel size and weight were reduced significantly. We assayed genetic differences between the two groups using selection statistics XP-EHH, XP-CLR, and F_ST_. All three selection statistics show evidence of selection at genomic regions associated with tassel morphology relative to genome-wide null distributions. These results support the tremendous effect, both phenotypic and genotypic, that selection has had on maize male inflorescence morphology.

## Introduction

The male inflorescence, or tassel, of maize (*Zea mays* L.) is a branched structure that displays remarkable morphological diversity. Tassels vary in length, number of branches, compactness, and curvature, all of which are capable of affecting the quantity and dispersion of pollen. Inflorescence architecture is crucial to reproduction in wind pollinated plants (Friedman and Barrett 2009) such as maize. As such, tassel size and shape are likely to have been subject to both natural and artificial selection. Tassels on teosinte plants, the wild progenitor of maize, are characterized by a large number (>50) of branches (Xu *et al*. 2017), implying a reproductive advantage of numerous branches in natural populations. In cultivated maize, however, tassel size and branch number have decreased over the course of modern breeding (Meghji *et al*. 1984; Duvick 1997, 2005). Tassel size is negatively correlated with grain yield, putatively due to shading caused by large tassels (Duncan *et al*. 1967; Hunter *et al*. 1969). At high planting densities typical of modern agronomic practices, tassels can intercept enough sunlight to reduce photosynthesis lower in the canopy, which can negatively impact assimilation of photosynthate in the ear (Duncan *et al*. 1967; Wardlaw 1990; Hammer *et al*. 2009). A study in tropical maize indicated that direct selection on tassel branch number can result in an indirect increase in grain yield (Fischer *et al*. 1987), showing that the relationship between tassel and ear may be causative and persists beyond temperate germplasm.

Previous studies have identified quantitative trait loci (QTL) and quantitative trait nucleotides (QTN) associated with different aspects of tassel morphology and development that colocalize with *teosinte branched1* (*tb1*) and *teosinte glume architecture1* (*tga1*), which were targets of domestication that changed plant and inflorescence architecture between teosinte and modern maize (Doebley *et al*. 1995; Wang *et al*. 2005; Brown *et al*. 2009; Wu *et al*. 2016). A study mapping tassel architecture in a population of recombinant inbred lines (RILs) derived from a cross between maize and teosinte identified a QTL and evidence of selection at the *barren infloresence2* (*bif2*) locus (Xu *et al*. 2017), which has been characterized previously as decreasing the number of tassel branches and spikelets (McSteen and Hake 2001). The same study identified five additional QTL that colocalized with inflorescence development genes and were in the list of genes identified as putatively selected during maize domestication by Hufford and et al. (2012). These findings provide evidence for selection on tassel morphology during domestication; however, tassel morphology has continued to change post-domestication, including significant differences observed within breeding germplasm developed between the early 20^th^ century and modern day (Meghji *et al*. 1984; Duvick 2005).

Until 1935, most of the maize grown in the North American Corn Belt consisted of open-pollinated populations, in which plants are allowed to reproduce freely with each other. Within four years, however, more than 90% of maize grown in Iowa had transitioned to higher-yielding hybrids created by controlled crosses, and the rest of the Corn Belt soon followed (Reif *et al*. 2005). The transition to single-cross hybrids was largely driven by key founder lines (B14, B37, and B73) derived from a pivotal maize population called Iowa Stiff Stalk Synthetic (BSSS), which was created in the 1930s by intermating 16 inbred lines (Lamkey *et al*. 1991; Reif *et al*. 2005). These founder lines and their combinations, referred to as Stiff Stalk lines, became preferentially used as females in hybrid breeding schemes, while other inbred lines that combined well with them were used as males (Reif *et al*. 2005). These early choices laid the framework for what would eventually become the heterotic groups used in maize breeding today. Descendants of the BSSS population have been used universally across proprietary breeding programs, resulting in a number of commercial Stiff Stalk lines protected by Plant Variety Protection certificates (PVP lines) (Mikel and Dudley 2006).

The shift in breeding methods to single-cross hybrids placed an emphasis on developing inbred lines with characteristics that enhanced their potential to serve as parents. These parents are subsequently crossed to each other to produce vigorous hybrid offspring. In open-pollinated populations, a competitive advantage was conferred by greater pollen production and dispersal (Friedman and Barrett 2009). Inbred line breeding greatly reduced that competitive advantage as breeders began making more controlled crosses; there is also evidence that inbreeding tends to shift resource allocation to the female inflorescence (Burd and Allen 1988). Thus, we hypothesized that selection in the Stiff Stalks by breeders between the original BSSS population and modern PVP lines acted to reduce overall tassel size.

Among the germplasm used in this study are inbred lines derived from the original BSSS population, as well as a number of commercially developed inbred lines for which the Plant Variety Protection certificates have recently expired (ex-PVPs). These ex-PVP lines represent the most modern germplasm from private seed companies that is available to the public, and they can be traced back to founder lines from the BSSS population. These two sets of materials afford us an opportunity to evaluate changes in maize tassel architecture caused by the accelerated evolutionary process of modern breeding. We used single nucleotide polymorphisms (SNPs) identified as significant in association mapping as *a priori* selection candidates, and compared selection statistics at those SNPs to their genome-wide distribution. If selection by breeders between the BSSS and modern ex-PVPs acted to reduce tassel size, we expect that loci associated with tassel morphology phenotypes would show enrichment for selection signatures relative to the rest of the genome. Indeed, we observe an enrichment of selection statistic scores in regions containing SNPs associated with tassel morphological traits, a finding that is consistent across three different selection statistics.

## Results

### Tassels exhibit morphological diversity for traits measured manually and by image-based methods

Manual measurements of tassel length (TL), spike length (SL), branch number (BN), branch density (BD), branch zone length (BZ), spike proportion (SP), and tassel weight (TW) for >22,000 individual tassels were recorded for two populations, an association panel (WiDiv-942; Mazaheri et. al, in press) and a nested association mapping population (PHW65 NAM) with PHW65 as the common parent and PHN11, MoG, and Mo44 as founder parents . These materials were grown in three environments between 2013 and 2015 in south central Wisconsin. The WiDiv-942 association panel used in this study is comprised of 942 diverse inbred lines, while the PHW65 NAM consists of three nested biparental populations of 200 individuals each. The WiDiv-942 was chosen to capture broad phenotypic and genetic diversity, whereas the PHW65 NAM leverages controlled biparental crosses between individuals with contrasting tassel morphologies. All tassels manually measured in 2015 were also photographed and measured using the image-based tassel phenotyping system TIPS (Gage *et al*. 2017), resulting in image-based measurements of TL, SL, BN, and TW (referred to as TLp, SLp, BNp, and TWp, respectively), along with four other traits: compactness (CP), fractal dimension (FD; a measure of overall complexity), skeleton length (SK; a measure of total linear length), and perimeter length (PR; the length of a tassel’s 2D outline).

All fifteen tassel morphological traits measured manually or by image-based phenotyping showed variability for raw measurements within both the WiDiv-942 and PHW65 NAM populations (Table 1; Supplemental Figure 1). Excluding FD, which ranged from 1.1 to 1.6, the other 14 traits exhibited between 2-fold (SLp) and 36-fold (BNp) variability within the PHW65 NAM and between 2-fold (SP) and 22-fold (SK) variability within the WiDiv-942.

**Table 1:** Summary of traits and heritabilities

| Trait Name (units) | Method | Abbreviation | Mean |  | Standard Deviation |  | Heritability / Repeatability |  |
| --- | --- | --- | --- | --- | --- | --- | --- | --- |
|  |  |  | PHW65<br>NAM | WiDiv-942 | PHW65<br>NAM | WiDiv-942 | PHW65<br>NAM | WiDiv-942 |
| **Tassel Length (cm)** | Manual | TL | 35.82 | 32.39 | 4.57 | 4.81 | 0.89 | 0.95 |
|  | Image | TLp | 35.83 | 34.21 | 3.36 | 3.99 | 0.63 | 0.79 |
| **Tassel Weight (g)** | Manual | TW | 16.11 | 15.6 | 7.16 | 6.86 | 0.94 | 0.96 |
|  | Image | TWp | 15.38 | 16.17 | 5.41 | 5.5 | 0.71 | 0.86 |
| **Branch Number (count)** | Manual | BN | 10.2 | 12.03 | 3.82 | 6.35 | 0.94 | 0.97 |
|  | Image | BNp | 11.1 | 12.17 | 3.81 | 3.74 | 0.70 | 0.82 |
| **Spike Length (cm)** | Manual | SL | 23.73 | 22.52 | 3.66 | 4.37 | 0.81 | 0.95 |
|  | Image | SLp | 25.36 | 22.77 | 2.62 | 3.1 | 0.63 | 0.79 |
| **Branch Zone (cm)** | Manual | BZ | 10.87 | 9.87 | 2.62 | 3.03 | 0.72 | 0.93 |
| **Spike Proportion (proportion)** | Manual | SP | 0.69 | 0.69 | 0.06 | 0.08 | 0.72 | 0.93 |
| **Branch Density (count/cm)** | Manual | BD | 0.97 | 1.21 | 0.32 | 0.51 | 0.80 | 0.93 |
| **Compactness (proportion)** | Image | CP | 0.34 | 0.34 | 0.11 | 0.14 | 0.70 | 0.76 |
| **Fractal Dimension (dimension)** | Image | FD | 1.42 | 1.42 | 0.04 | 0.05 | 0.67 | 0.81 |
| **Perimeter Length (pixels)** | Image | PR | 10902.52 | 10399.72 | 3167.7 | 3585.34 | 0.68 | 0.72 |
| **Skeleton Length (pixels)** | Image | SK | 5680.85 | 5585.8 | 2021.27 | 2377.85 | 0.72 | 0.78 |

Descriptive statistics of fifteen tassel morphological traits measured in two populations. Image-based traits are highlighted in grey, manually measured traits in white. All manual traits except Tassel Weight were measured in 3 environments, while Tassel Weight and all image traits were measured in one environment. Means and standard deviations were calculated from untransformed per-plot averages of measurements taken on up to three plants.

The 15 traits can be broadly classified into two groups: those that contribute to tassel size by contributing to length (TL, TLp, SL, SLp, SP, BZ), and those that contribute to tassel size by contributing to “branchiness” (BN, BNp, BD, CP, FD, SK, PR, TW, TWp). Though TW and TWp are measures of mass rather than branching, both traits are affected more by changes in branchiness than changes in tassel length and therefore are included in the former group.

For most traits, the PHW65 NAM has a higher or equivalent mean to the WiDiv-942, with the exception of TWp (0.79 grams lighter), BN (1.83 fewer branches), BNp (1.07 fewer branches), and BD (0.24 fewer branches/cm). As expected given the allelic representation contained in each of the populations, phenotypic variability within the PHW65 NAM is lower than in the WiDiv-942, with the exception of TW (standard deviation 0.3 grams greater) and BNp (standard deviation 0.07 branches greater).

Heritabilities for manually measured traits ranged from 0.93 (BZ) to 0.97 (BN) in the WiDiv-942 and from 0.79 (BZ) to 0.94 (BN) in the PHW65 NAM. Since images and TW were taken only at one location, repeatability was calculated for image-based traits and TW in place of heritability. Repeatabilities ranged from 0.72 (PR) to 0.86 (TWp) in the WiDiv-942 and 0.63 (TLp) to 0.72 (SK) in the PHW65 NAM (Table 1). The high heritabilities for measurements taken in multiple environments indicate large genotypic variance relative to genotype-by-environment variance and error. Similarly, high repeatabilities for traits that were measured in a single environment indicate large genotypic variance relative to error. It is unsurprising that the image-based repeatabilities were lower than the manually measured heritabilities, as manual measurements were taken in three separate environments while image-based traits were measured in one environment.

### Tassel morphology has changed over the course of 20^th^ century maize breeding

The WiDiv-942 association panel contains 41 inbred lines developed by selfing without selection from cycle 0 of the BSSS population (BSSSC0), 16 public inbred lines with pedigrees that trace entirely back to lines derived from the BSSS, and 21 ex-PVPs with genotypic evidence for descent from the BSSS (Supplemental File 1). These three groups of lines allow comparison of tassel morphology over the course of modern maize breeding. The BSSSC0 lines represent the earliest material, the ex-PVPs represent the most recent and heavily selected material, while the public lines form an intermediate group.

Comparison of best linear unbiased predictors (BLUPs) for each group revealed that the mean values of public inbred BLUPs for each trait showed between a 2% (FD, TLp) and a 16% (TWp) change relative to the BSSSC0, while ex-PVP lines displayed between a 4% (TLp) and a 42% (TWp) change in mean relative to the BSSSC0 lines (Figure 1). For all traits except CP, the public inbred lines displayed intermediate phenotypes between the medians of the BSSSC0 and ex-PVP lines. Public inbred lines were significantly different from BSSSC0 lines for BN, BD, FD, TWp, and BNp, while ex-PVP lines were significantly different from BSSSC0 lines for all traits except TL, TLp, and CP. Public lines and ex-PVP lines differed significantly for BN, SP, BD, CP, FD, TW, TWp, and BNp (all comparisons: Tukey’s HSD, α = 0.05). Generally, tassel length (TL and TLp), spike length (SL and SLp), and spike proportion (SP) increased over time, while all other traits decreased over time. The observed phenotypic trends suggest selection on tassel morphology between the open-pollinated BSSS and public inbreds, as well as continued selection between the public lines and the ex-PVP lines.

**Figure 1:**
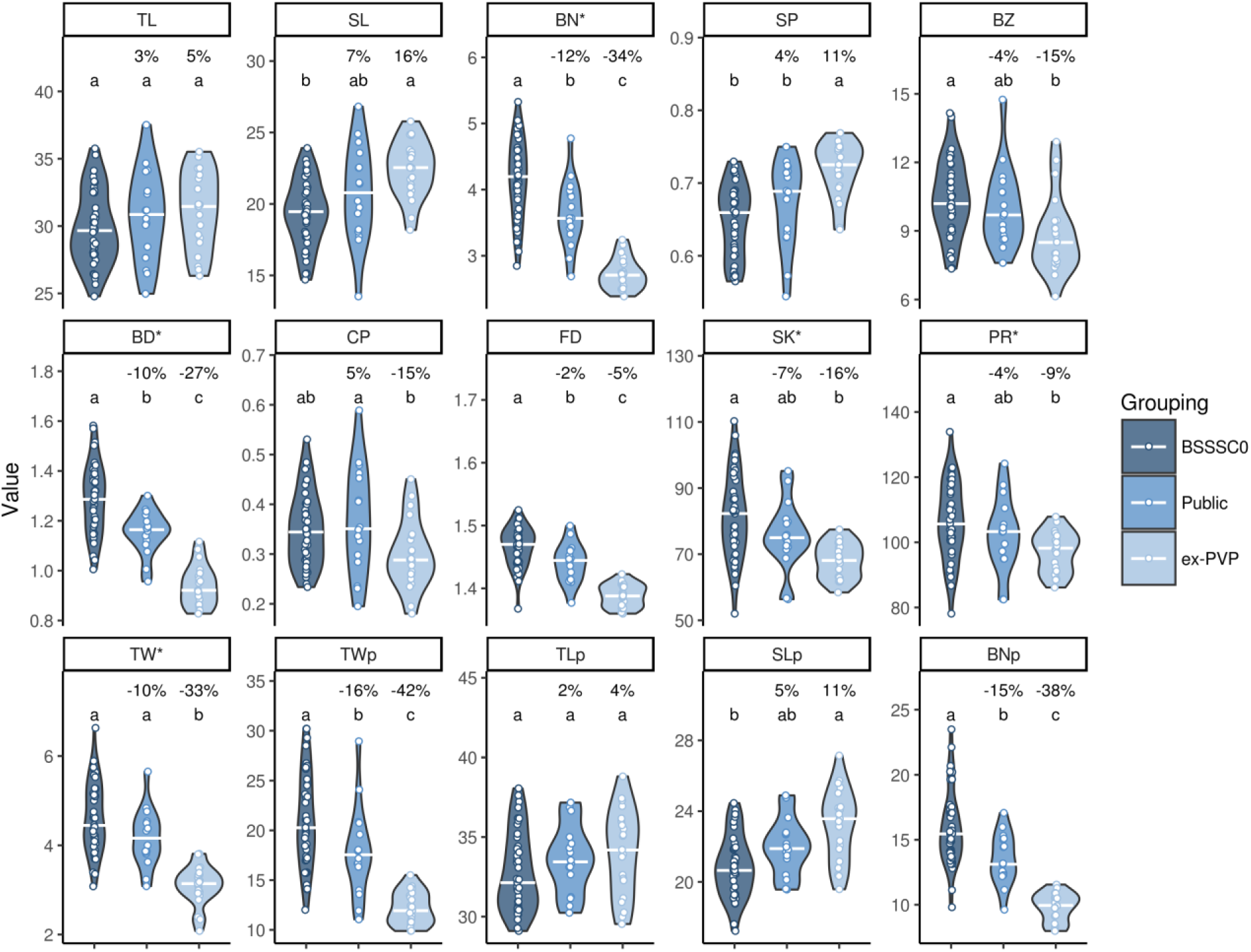
Tassel traits show progressive changes over time. Best linear unbiased predictors (BLUPs) of 15 tassel morphological traits for 41 BSSSC0 inbreds, 16 publicly released inbreds derived from BSSS lines, and 21 ex-PVP inbreds derived from BSSS lines. Percentages indicate the percent change in mean value relative to the BSSSC0 lines. White bars indicate the median value. Letters indicate significant differences between groups (Tukey’s honest significant difference test, α = 0.05). Trait abbreviations noted with an asterisk were square root transformed before computing BLUPs. TL: tassel length; SL: spike length; BN: branch number; SP: spike proportion; BZ: branch zone; BD: branch density; CP: compactness; FD: fractal dimension; SK; skeleton length; PR: perimeter length; TW: tassel weight; TWp, TLp, SLp, and BNp: image-based predictions of TW, TL, SL, and BN.

### GWAS identifies SNPs associated with tassel morphological traits

All 15 traits were mapped in the WiDiv-942 association panel using the multiple locus linear mixed model implemented in FarmCPU (Liu *et al*. 2016) with 529,018 single nucleotide polymorphisms (SNPs) discovered from RNA sequencing with minor allele frequency >0.02. Traits were mapped in the PHW65 NAM populations using a resampling-based GWAS approach (Tian *et al*. 2011) with 10.6 million resequencing SNPs projected from parental lines onto the progeny, which were genotyped at 9,291 genotyping-by-sequencing (GBS) SNPs. In the WiDiv-942, 87 SNPs were identified as significantly associated with 12 of the 15 traits. In the PHW65 NAM, 155 SNPs were identified as significantly associated across all traits (Supplemental File 2).

The manually measured traits returned a greater number of significant associations in total than the image-based traits, as expected based on the relative measures of heritability and repeatability. In ten instances, manual and image-based measures of the same phenotype were associated with the same SNP: one SNP for tassel length (TL and TLp), four for branch number (BN and BNp), one for spike length (SL and SLp), and five for weight (TW and TWp). One pair of SNPs each for branch number and weight were identified in the WiDiv-942, and the rest of the colocalized SNPs were discovered in the PHW65 NAM. The single SNP associated with tassel length was the same as the single SNP associated with spike length; a SNP associated with weight and a SNP associated with branch number were within 2.5kb of each other. In two other instances, the manually measured and image-based traits were associated with SNPs within 5kb of each other, and in an additional two instances they were associated with SNPs that were located within 150kb of each other (Supplemental Figure 2). Generally, there is more concordance between SNPs associated with manual and image-based measurements for traits with higher heritabilities/repeatabilities (branch number and tassel weight) than lower heritabilities/repeatabilities (spike and tassel length). This is to be expected, as genetic signal is stronger when heritability is high, but the pattern seen in this study may be due only to chance as BN and TW are only marginally more heritable than spike and tassel length.

### Selection statistics show evidence for selection at SNPs associated with tassel morphology

To test for genomic signatures of selection on tassel morphology, SNPs associated with tassel characteristics in the WiDiv-942 and PHW65 NAM were used as *a-priori* candidates for regions having undergone selection between the BSSSC0 lines and the ex-PVP lines. Scans for selection were conducted using linkage disequilibrium (LD) focused methods XP-EHH (Sabeti *et al*. 2007) and XP-CLR (Chen *et al*. 2010), and the allelic frequency change based method F_ST_ (Weir and Cockerham 1984). Genome-wide results from selection scans were binned into 10kb windows, and windows containing SNPs associated with tassel morphology were compared to the genome-wide distribution of each statistic. Of the 242 total SNP-trait associations identified, 170 were in windows that had XP-EHH scores, 231 had XP-CLR scores, and 169 had F_ST_ scores. Not all SNP-trait associations had scores for each selection statistic because the associations were discovered in the full PHW65 NAM and WiDiv-942 populations, but may lie in regions that are monomorphic between the BSSSC0 lines and ex-PVP individuals or contain missing SNP calls. The distributions of tassel-associated XP-EHH, XP-CLR, and F_ST_ scores were significantly different from the genome-wide permuted null distributions (Kolmogorov-Smirnov test; p = 1.8 × 10^-3^, 7.8 × 10^-3^, and 1.9 × 10^-3^, respectively), suggesting enrichment for selection statistics in regions associated with tassel morphology.

XP-EHH scores for tassel-associated windows (Figure 2) were enriched for values approximately greater than 1 and less than −1, with the majority of the enrichment coming from negative scores (Figure 3). Positive XP-EHH scores indicate longer, more prevalent haplotypes in the collection of lines from the BSSSC0 population, while negative XP-EHH scores indicate longer, more prevalent haplotypes in the ex-PVP set. The greater surplus of negative scores among the tassel-associated SNPs indicates extended haplotypes at those loci in the newer ex-PVP population, relative to the ancestral BSSS. The extended haplotypes in the ex-PVP population are consistent with what would be expected if beneficial alleles were recently selected, resulting in a concurrent increase in linked haplotypes due to genetic hitchhiking (Voight *et al*. 2006). Further, the excess of negative XP-EHH scores was driven largely by associations with phenotypes related to branchiness rather than length (Figure 4; One-sided Wilcoxon rank sum test, p=2.9 × 10^-3^). Branchiness traits have more impact on tassel size and biomass than tassel and spike length. With the exception of BZ, all the length traits increased from the BSSSC0 to the ex-PVPs, whereas all the branchiness traits decreased. As such, most of the SNPs showing evidence of selection in the ex-PVPs are associated with traits that control overall tassel size and biomass and have decreased significantly since the 1930s.

**Figure 2:**
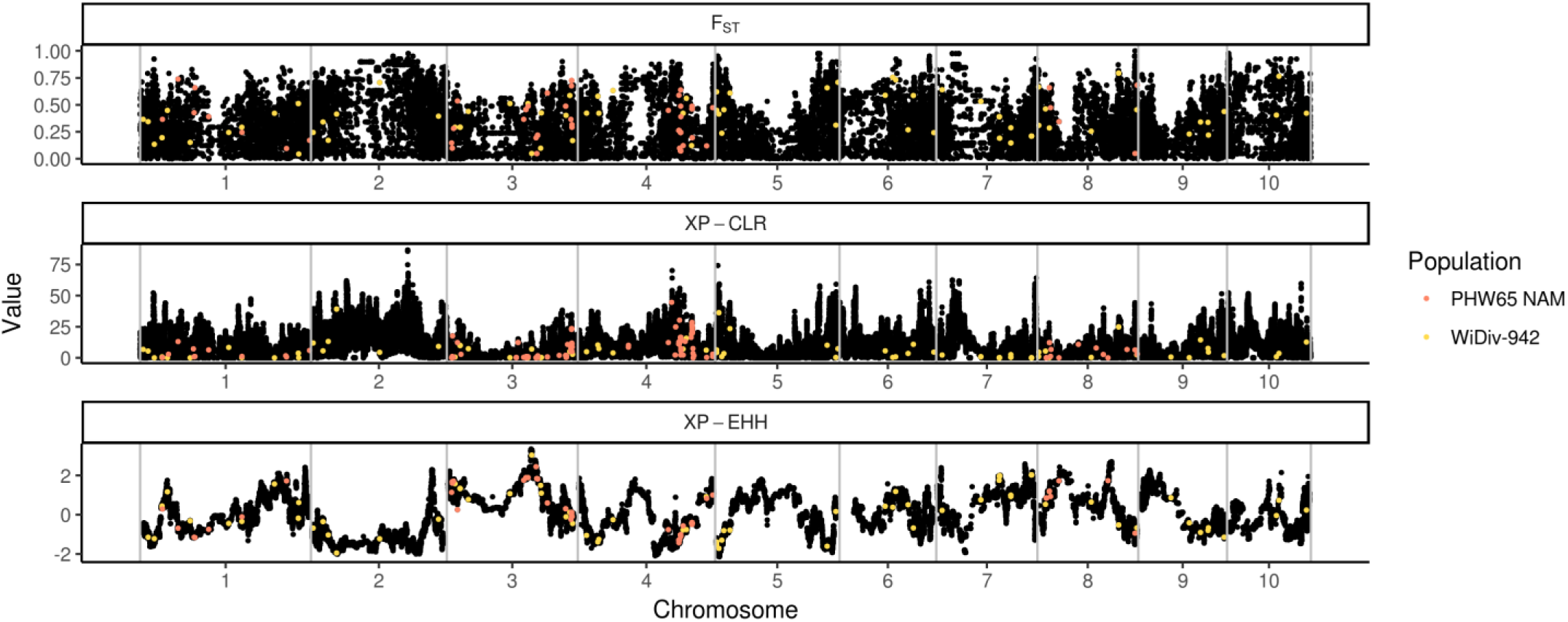
Physical locations and selection statistics for SNPs associated with tassel morphological traits (Genomwide_statistics.png) Values of maximum F_ST_ (top), XP-CLR (middle), and XP-EHH (bottom) in nonoverlapping10kb windows along the genome (black). Windows containing single nucleotide polymorphisms associated with tassel traits are highlighted in color; those in yellow were discovered by GWAS in the WiDiv-942 association panel, while those in red were discovered in the PHW65 NAM.

**Figure 3:**
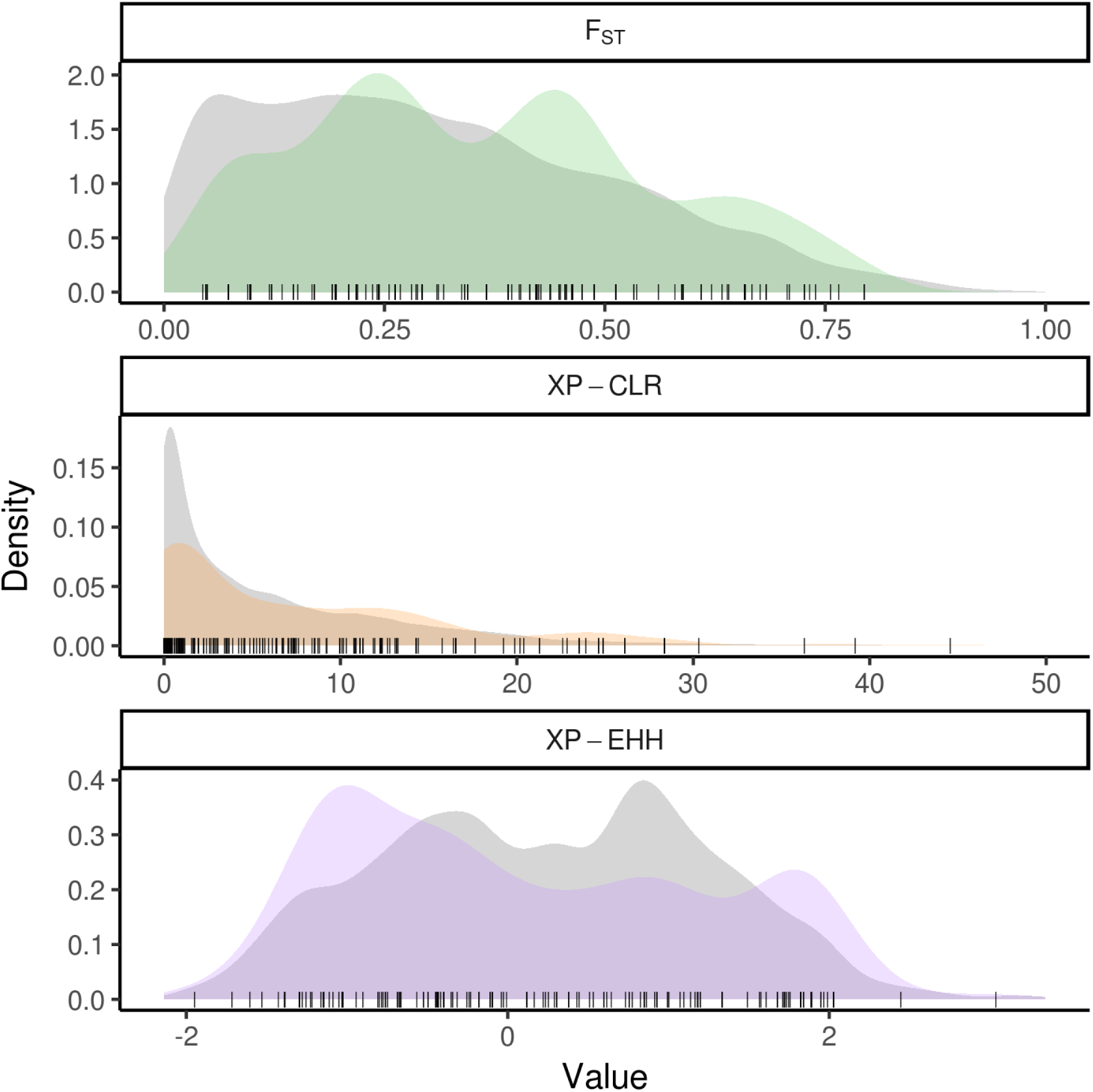
Densities of selection statistics for SNPs associated with tassel morphological traits reveal enrichment for signals of selection (Distributions.png) Distributions of selection statistics for single nucleotide polymorphisms (SNPs) associated with tassel morphological traits (colored), compared to genome-wide null distributions (grey) derived by circular permutation. Values of individual SNPs associated with tassel traits are represented by points along the bottom of each plot. The XP-CLR plot was truncated at 50 for better visualization.

**Figure 4:**
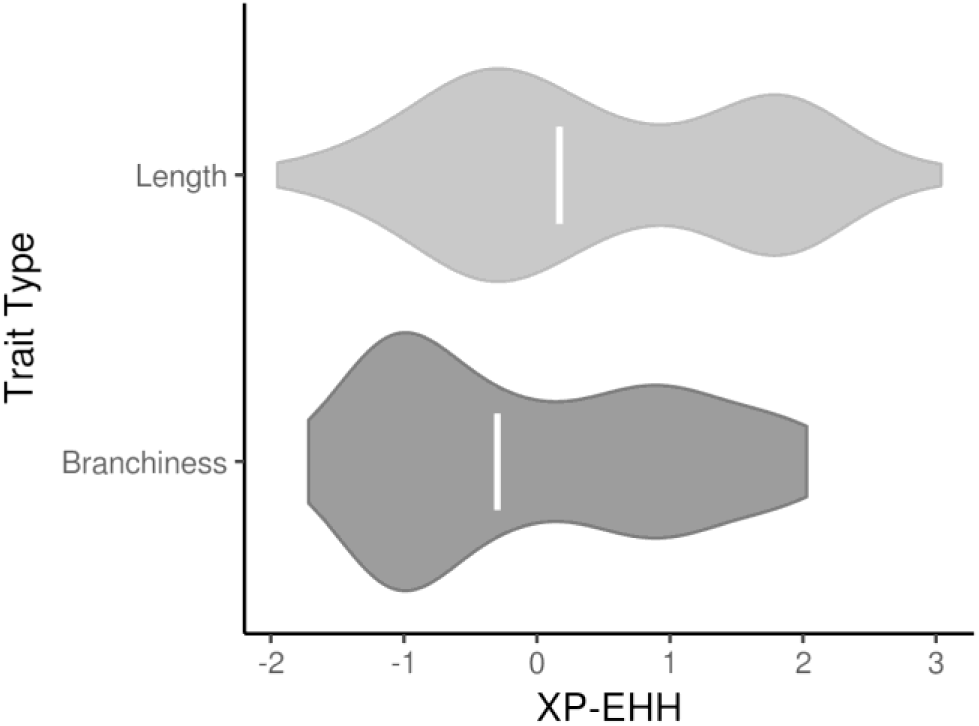
Signals of selection for tassel traits are driven by traits related to quantity and density of branches (XPEHH_by_trait_type.png) Distribution of XP-EHH values for tassel trait associated SNPs, categorized into SNPs associated with branchiness traits (n=141) and those associated with length traits (n=88). The enrichment of negative XP-EHH values, suggestive of selection in the ex-PVP lines, is driven by traits related to branchiness. White bars indicate median value.

Similar to XP-EHH, XP-CLR is a haplotype-based method for identifying selective sweeps. Rather than identify alleles that have recently arisen to substantial frequency, as XP-EHH does, XP-CLR identifies alleles that have changed in frequency more quickly than expected under neutrality (Chen *et al*. 2010). Both methods use LD around the tested allele as a measure of the allele’s age. Tassel-associated XP-CLR scores (Figure 2) show enrichment for values in the tail of the distribution (Figure 3), indicating an excess of regions that may have been subject to selective sweeps.

F_ST_, a measure of population differentiation based on allele frequencies, differs from the previous two selection statistics in that it is not haplotype based. F_ST_ values for tassel-associated windows (Figure 2) are generally higher than expected under the null distribution, with a depletion of values less than approximately 0.3 and enrichment for values greater than approximately 0.3 (Figure 3).

For the 101 significant associations that were genotyped in the WiDiv-942 (and hence, in the ex-PVP and BSSSC0 lines), we hypothesized that alleles with higher frequency in the ex-PVPs would generally have allelic effects conferring more ex-PVP-like phenotypes, that is, positive values for TL, SL, TLp, SLp, and SP, and negative values for all other traits. However, we found no difference between the effect sizes of alleles with higher frequency in the ex-PVPs and alleles with higher frequency in the BSSSC0s (Welch’s two-sampled t-test, p=0.87). The lack of any difference could be due to the low number of SNPs we were able to directly test, or perhaps due to noise introduced by identifying SNPs that are not causative themselves but in LD with variants that directly affect tassel morphology.

None of these three selection statistics show strong, systematic relationships with each other on a genome-wide basis (Supplemental Figure 3) or among the tassel-associated windows (Supplemental Figure 4). However, there are two tassel-associated windows with XP-EHH less than −1.6 and XP-CLR greater than 35, located near the beginning of chromosome 2 (46.11-46.12Mb) and the beginning of chromosome 5 (6.79-6.80Mb). Both windows have F_ST_ scores of 0.4. The significant SNP on chromosome 2 is located 7bp downstream of the gene Zm00001d003499, which is annotated as a RING/U-box superfamily protein. The *Arabidopsis thaliana* gene BIG BROTHER is a RING/U-box superfamily protein encoding gene that has been shown to negatively regulate floral organ size (Disch *et al*. 2006), suggesting that variability in Zm00001d003499 may have been selected on between the BSSS and ex-PVP lines and also be involved in inflorescence morphology.

## Discussion

Tassel morphology of modern commercial inbred lines, which are used as parents of single cross hybrids grown in the United States Corn Belt, is dramatically different from tassel morphology seen in the open-pollinated varieties of the pre-hybrid era. We demonstrate that tassel morphology has been continually modified, with length-related traits increasing and branchiness traits decreasing. This trend is consistent between BSSSC0 inbreds and public BSSS-derived inbreds, as well as between public inbreds and BSSS-derived ex-PVPs. Through mapping of both image-based and manually measured phenotypes, we identify SNPs associated with tassel morphological traits, which collectively show enrichment for three selection statistics.

By studying traditionally measured traits, such as tassel length and branch number, as well as traits that cannot be accurately measured manually, such as compactness and fractal dimension, we generate a more comprehensive and nuanced representation of tassel morphology than in previous tassel studies. Previous QTL mapping and GWAS of maize tassel morphology have primarily emphasized tassel branching and length (e.g., Berke and Rocheford 1999; Mickelson *et al*. 2002; Upadyayula *et al*. 2006; Brown *et al*. 2011; Wu *et al*. 2016). The novel, image-based traits used in this study display considerable variability, have high enough repeatabilities to be useful in mapping studies, and resulted in significant phenotype-genotype associations that contributed to our analysis of selection signals. The ability to add information to manual phenotyping is especially impactful when manual phenotyping is laborious or inefficient. For phenotyping tassels, manual measurements are relatively easy and fast to obtain. However, image-based phenotyping measures the same traits but also contributes, with no additional time investment, additional measurements such as compactness and fractal dimension, which would be difficult, subjective, or impossible to measure otherwise. Though tassels for this study were removed from the plants before imaging, a laborious task, such methods are a step towards higher throughput methodologies, such as image acquisition by unmanned aerial vehicles.

By comparing tassel morphology in the BSSSC0 and ex-PVP sets considered in this study, we are able to compare two groups that bookend modern breeding of one of the most important heterotic groups in North American maize germplasm. The BSSSC0 and ex-PVP groups are characterized by unique and differing tassel morphology: relative to the BSSSC0, ex-PVP tassels are longer, less branched, and weigh less. These changes in morphology reflect an overall trend towards a tassel ideotype that has few lateral branches and produces pollen primarily by means of an extended central spike. The Stiff Stalks, traditionally used as females in breeding programs, need only produce enough pollen for self-fertilization during inbred line development and seed increases; this is reflected by the reduction of their size and weight since the inception of inbred-driven breeding schemes in the 1930s. Previous experiments that recurrently selected the BSSS population for grain yield also observed a decrease in tassel branch number (Edwards 2011), further reinforcing the relationship between agronomic performance and reduced tassel size. Though the changes in tassel morphology caused by recent selection are well characterized, the genetic effects of selection by breeders are less studied.

Previous works have shown evidence for regions selected during domestication in animals such as pigs and chickens (Rubin *et al*. 2010; Yang *et al*. 2014); and in plants such as maize and soybean (Hufford *et al*. 2012; Zhou *et al*. 2015). Evidence has been found for differential artificial selection between breeds of cattle (Rothammer *et al*. 2013) as well as for post-domestication improvement in maize and soybean (Hufford *et al*. 2012; Zhou *et al*. 2015), but the latter two characterize selection between landraces and improved varieties, which cover a much larger period of selection than just modern breeding. Work in rice (Xie *et al*. 2015) has identified overlap between inflorescence morphology QTL and regions showing evidence of selection in the last 50 years of breeding, but the genetic relationship between recent selection and inflorescence morphology in maize is relatively unexplored.

XP-EHH, XP-CLR, and F_ST_ scores for SNPs associated with tassel morphology traits showed deviations from permuted genome-wide null distributions, with enrichment for values that are indicative of selection. In *de novo* scans for selected sites, often the threshold for determining selection candidates by XP-CLR or F_ST_ is set as a quantile of the empirical distribution that limits candidates to loci in the top, e.g., 0.1% or 0.01% of results (e.g., Beissinger *et al*. 2014; Jeong *et al*. 2015). XP-EHH has the agreeable property of being approximately normally distributed under a model of no selection, which allows identification of selection candidates based on p-values from the normal distribution (Sabeti *et al*. 2007). While not all tassel-associated SNPs have high enough XP-EHH, XP-CLR, and F_ST_ scores to be identified as selection candidates in a *de novo* search for selected sites, the strong enrichment for elevated scores suggests identification of regions that might be experiencing weak or ongoing selection. Genome-wide scans for selection are complicated by population dynamics and demographics that differ from expectations, and often result in identification of large genomic regions containing many features that require further investigation. Instead of approaching such an analysis with no expectations, it has been suggested (Haasl and Payseur 2016) that creating an *a priori* list of candidate genes can help narrow down significant results. In this study, we used associations from two comprehensive genetic mapping resources as *a priori* candidates to identify signatures of selection associated with a specific suite of phenotypic traits.

XP-EHH identifies putatively selected sites by identifying regions where one population contains longer haplotypes around a target SNP, which have arisen due to hitchhiking with a selected locus. We observe enrichment for both positive and negative XP-EHH scores, which indicate greater haplotype homozygosity in the BSSSC0 and ex-PVP sets, respectively. There is a substantially larger enrichment for negative XP-EHH scores than positive, consistent with the hypothesis that favorable alleles from the BSSSC0 lines were selected in the ex-PVPs, resulting in increased LD in the surrounding areas. The majority of enrichment for negative XP-EHH scores appears to be driven by “branchiness” traits, which may have been easier to select on (intentionally or not) by breeders aiming to reduce overall tassel size. The enrichment for positive XP-EHH scores, though not as extreme as for negative scores, indicates a number of tassel-associated loci with slower LD decay in the BSSSC0 lines than in the ex-PVP set. One possible explanation for this observation is that prior to the creation of the BSSS, during a period characterized by open-pollinated populations, heavier tassels with more branches had a reproductive advantage due to higher pollen production. During the last century of maize breeding the selective pressure on high pollen production has been removed by a hybrid breeding scheme in which inbred combining ability and hybrid grain yield are prioritized. It is possible that the BSSSC0 lines show extended haplotypes due to selection favoring alleles that confer large tassel size, whereas modern breeding schemes have removed that selective pressure and allowed or encouraged recombination in previously selected regions.

The results presented here provide evidence for selection by 20^th^ century maize breeders on regions associated with maize tassel morphology. Whether these changes are the result of direct or indirect selection on tassel size and shape is unclear. Modern, commercially developed maize inbreds display a suite of characteristics that have systematically changed since the open-pollinated varieties of the 1930s, and it is difficult to disentangle selection for changes in tassel morphology from selection for traits that are regulated by loci pleiotropically involved in tassel morphology. Because the tassel and the ear follow a shared developmental path early in development, genes affecting morphology of one may very well have effects on the other. QTL have been discovered that exhibit pleiotropic behavior, conferring a positive relationship between tassel and ear length (Brown *et al*. 2011). Mutants of the inflorescence genes *unbranched2* and *unbranched3* have been shown to simultaneously decrease branch number and increase kernel row number, a trait that directly affects grain yield (Chuck *et al*. 2014), while mutants of *zea floricaula/leafy 2* decrease both branch number and kernel row number (Bomblies and Doebley 2006). There is also evidence for shared genetic control of tassel traits and other plant architecture traits such as leaf length (Brown *et al*. 2011) and leaf angle (Mickelson *et al*. 2002), the latter of which has also changed significantly over the course of modern maize breeding (Duvick 2005). Though more work is needed to clarify the genetic relationships between selection and maize tassel morphology, particularly with respect to disentangling pleiotropic effects, this study is a starting point for discovering the effects artificial selection has on genetic control of inflorescences in agricultural crops.

Change in tassel size and shape between the early 20^th^ century and modern day is a well-characterized phenomenon but the causes of that phenomenon and its potential causal effects on other agronomic traits and productivity are not fully known. Our results present evidence for enriched signals of selection in genomic regions associated with tassel morphological traits. Further, this selection took place in approximately 60 years between the creation of the BSSS in the 1930s and the release of ex-PVP lines in the 1980s and 1990s, a tremendous example of the efficiency of modern crop breeding in imposing drastic genetic and phenotypic changes.

## Materials and Methods

### Populations

This study used two populations of maize inbred lines: a diversity panel and a nested association mapping population. The diversity panel (WiDiv-942) consists of 942 diverse lines that typically reach grain physiological maturity in the Midwest regions of the United States, and represents an expansion of the 503 line Wisconsin Diversity panel (WiDiv-503; Hirsch *et al*. 2014) to increase breadth and depth of genetic diversity as well as to include a greater representation of ex-PVP (lines formerly protected by Plant Variety Protection certificates) inbreds. The nested mapping population (PHW65 NAM) was styled as a smaller version of the maize NAM (McMullen *et al*. 2009) and consists of three biparental populations that share a common parent, the inbred line PHW65. PHW65 is an ex-PVP Lancaster-type line. The three founder lines are PHN11, Mo44, and MoG. PHN11 is an ex-PVP Iodent line closely related to PH207; Mo44 is a Missouri line derived from Mo22 and Pioneer Mexican Synthetic 17 (Flint-Garcia *et al*. 2005); and MoG is a Missouri line characterized by wide leaves, a heavy stalk, and high pollen production. The founder lines were chosen for their differing tassel morphologies: MoG and Mo44 have longer tassels than the recurrent parent PHW65; PHN11 has more branches than PHW65; the MoG tassels are heavy and curved while the PHN11 tassels are light and more rigid. Together, the four parental lines comprise a diversity of tassel size and shape. Each biparental population consisted of 200 individuals derived by double haploid generation from the F2 of the original parents.

The WiDiv-942 was grown in the summers of 2013 and 2014 at the University of Wisconsin’s Arlington Agricultural Research Station and in the summer of 2015 at the West Madison Agricultural Research Station. The PHW65 NAM was grown in 2014 at the Arlington Agricultural Research Station and in 2014 and 2015 at the West Madison Agricultural Research Station. All experiments were planted as randomized complete block designs with two replications at each location.

### Genotypic data

The WiDiv-942 lines were genotyped by RNA sequencing as described in Hirsch *et al*. (2014). RNA sequencing data were combined with RNA sequencing data from the original Hirsch *et al*. (2014) publication and aligned to the B73 version 4 reference genome sequence (Jiao *et al*. 2017) for SNP calling at 899,784 SNPs. The WiDiv-942 SNP matrix contained 30% missing data, which was imputed using fastPHASE v1.4.0 (Scheet and Stephens 2006) with the – H flag set to −3 to prevent haplotype phasing and default parameters otherwise.

Inbred lines in the PHW65 NAM population were genotyped using genotyping-by-sequencing (GBS) as described in (Elshire *et al*. 2011), using the GBSv2 pipeline implemented in TASSEL5 (Bradbury *et al*. 2007), resulting in 9,291 SNPs which were imputed with FSFHap (Swarts *et al*. 2014) using the “cluster” algorithm with a minimum MAF of 0.3 within each biparental population, a window size/step size of 30/15 for chromosomes 1, 4, and 6, and 50/25 otherwise.

The four parents of the PHW65 NAM population were resequenced at between 10x and 15x coverage and genotyped at 23.3 million SNPs (Brohammer *et al*. 2018). The 31% missing SNP data in the parents was imputed using fastPHASE v1.4.0 (Scheet and Stephens 2006) with the –H flag set to −3 to prevent haplotype phasing, the –K flag set to 7, and the –T flag set to 10. The –K flag sets the number of haplotype clusters at 7 and was chosen by evaluating imputation accuracy for K values between 1 and 31 and choosing the value above which minimal gains in accuracy were observed, similar to (Tian *et al*. 2011). The –T flag sets the number of random starts of the algorithm. As opposed to the imputation of the WiDiv-942, setting the –K and –T flags helped reduce the computational time of imputation for the larger number of resequencing SNPs. SNP calls from the four resequenced parents at 10.6 million polymorphic SNPs were projected onto their biparental progeny in the PHW65 NAM using GBS markers as a scaffold, as described in (Tian *et al*. 2011).

### Phenotypic measurements

Both populations were measured for tassel branch number (BN), tassel length (TL), and spike length (SL) on three representative plants per plot during or after flowering. BN was quantified as the number of primary tassel branches, TL was measured as the distance in cm from the lowest tassel branch to the tip of the spike, and SL was measured as the distance in cm from the uppermost tassel branch to the tip of the spike. All traits were measured in all environments with the exception of SL, which was not measured in the PHW65 NAM at West Madison in 2014. Three derived tassel traits can be calculated from these measurements: branch zone (BZ) as TL minus SL, branch density (BD) as BN divided by BZ, and spike proportion as SL divided by TL.

In addition to manual measurements, tassels measured in 2015 were also imaged and measured automatically using TIPS (Gage *et al*. 2017), which estimates TL and BN, as well as tortuosity (TR), compactness (CP), fractal dimension (FD), skeleton length (SK), perimeter length (PR), and tassel area (TA) from two-dimensional profile images of tassels. TR is a measure of tassel curvature expressed as a proportion: the length of a straight line from base to tip divided by the length of a spline fit along the tassel itself from base to tip; TA is the number of pixels in the tassel; CP is measured as TA divided by the area of the tassel’s convex hull; FD is a measure of overall tassel complexity; SK is a measure of overall linear length of the tassel, including the main spike and all branches; and PR measures the length of the tassel’s outline. Tassel weight (TW) was also measured as the mean dry weight of three tassels per plot.

Rather than directly use TA and the estimates of TL and BN produced by TIPS, we used partial lease squares regression (PLSR) models to generate more accurate, image-based predictions of TL, BN, SL, and TW. Models were fit using the *pls* package (Mevik and Wehrens 2007) in R (R Core Team 2016). Individual tassel measurements were used for model fitting and prediction was done using all eight model components (TIPS produces eight tassel measurements). The TIPS output for the PHW65 NAM population were used as explanatory variables and the model was trained on manually measured values of TL, SL, BN, and TW in the PHW65 NAM. The resulting PLSR models were used to predict image-based values of TL, SL, BN, and TW from the TIPS output for images of the WiDiv-942 population. The opposite was done as well – TIPS output for the WiDiv-942 population were used as explanatory variables in conjunction with manual measurements of TL, SL, BN, and TW to train the models, which were then used to predict image-based values of TL, SL, BN, and TW for the PHW65 NAM individuals. Image-based predictions of TL, SL, BN, and TW are referred to as TLp, SLp, BNp, and TWp.

The WiDiv-942 inbred line best linear unbiased predictions (BLUPs) for all manually measured traits, except TW, were calculated as the random genotypic effect predictions from the linear model 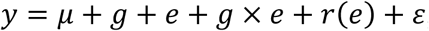, where *y* is the phenotypic response, *g* is the genotypic effect of an inbred line, *e* is the environmental effect of a year-location combination, (*g*×*e*) is the genotype-by-environment interaction, and *r*(*e*) is the effect of replicate within environment. The PHW65 NAM BLUPs for manually measured traits, except TW, were calculated from the model 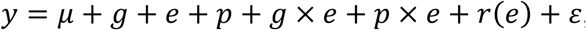, where *p* is an effect attributed to each biparental population, *p*×*e* is a population-by-environment effect, and all other terms are the same as in the WiDiv-942 model. BLUPs for image-based traits and TW for both populations were calculated from models identical to those above, but with all terms containing an environmental effect removed and a standalone replication term included. All effects were considered random, and models were fit with the R package *lme4* (Bates *et al*. 2016). BN, BD, PR, SK, and TW were square root transformed to better meet the model assumption of normally distributed residuals, and TR was removed from further analysis due to violation of model assumptions. All BLUPs can be found in Supplemental File 3.

Heritability was calculated on an entry-mean basis for all manually measured traits except for TW as 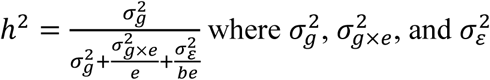 were estimated from the linear models described above and *b* and *e* are the number of replications and the number of environments, respectively. Repeatability was calculated similarly for all image-based traits and TW as 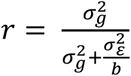, because image-based traits and TW were measured only in a single environment.

### Classification of public and ex-PVP stiff stalk lines

We identified lines that represent early Stiff Stalk germplasm, recent ex-PVP Stiff Stalk germplasm, and publicly developed Stiff Stalk germplasm that represents a temporally intermediate group between the two.

The WiDiv-942 contains 47 inbreds derived through self pollination from the original BSSS population (referred to hereafter as BSSSC0), which represent maize stiff stalk germplasm from the early 20^th^ century.

From all ex-PVP lines in the WiDiv-942, inbreds were chosen that showed genotypic evidence of being directly descended from the BSSS population (Supplemental File 1). Though pedigree information was available for some ex-PVPs, the number of individuals that could be traced by pedigree back to BSSS-derived inbreds was low. Additionally, using genotypic information to identify BSSS-derived ex-PVP lines allows exclusion of individuals showing evidence of contamination from other breeding material. Subpopulation structure within all the ex-PVP lines was estimated using ADMIXTURE (Alexander *et al*. 2009). The genotypic matrix consisting of 899,784 SNPs for 328 ex-PVP inbreds was first pruned for linkage disequilibrium using PLINK (Chang *et al*. 2015). Using a sliding window of 50 SNPs and a step size of 10 SNPs, SNPs with a R^2^ value greater than 0.1 within the window were discarded, leaving 120,000 SNPs. The pruned marker set was used in ADMIXTURE without seeding founding individuals. 10-fold cross validation of the errors was minimized at K groups equal to eight as an appropriate model for this marker set. These eight groups represent the resulting subpopulations; B14, B37, B73, Oh43, Lancaster, Iodent, Iodent-Lancaster, and Flint. To identify lines that both display evidence of descent from the BSSS as well as exclude lines with evidence of introgression from other germplasm pools, individuals were required to have a sum of >95% identity to the B14, B37, and B73 groups, as well as <2% identity to each of the other groups. Of the lines that met this requirement, 21 had been phenotyped and were kept for subsequent analysis. Those BSSS-descended ex-PVP lines are referred to hereafter as simply ex-PVPs. Hierarchical clustering of the ex-PVPs and BSSSC0 lines revealed six of the BSSSC0 lines did not cluster with the others. Those six were removed from further analysis, leaving 41 BSSSC0 lines.

The third group, referred to as “Public”, is comprised of 16 inbred lines from the WiDiv-942 that are known by pedigree to have been developed from the original Stiff Stalk synthetic population.

BLUPs for each of the three groups were compared for each of the 15 tassel morphological traits, and significant differences between BSSSC0, Public, and ex-PVP lines were evaluated by Tukey’s HSD (Tukey 1949).

### Mapping

Genome-wide association for the WiDiv-942 was performed in R (R Core Team 2016) using the multiple locus linear mixed model (MLMM) implemented in *farmCPU* (Liu *et al*. 2016). The first five principal components were included to account for spurious associations due to population structure, and the threshold for initial inclusion of SNPs into the model was set to allow an empirical false entry rate of 1% based on 1,000 permutations. SNPs with a minor allele frequency less than 0.02 were excluded from analysis, leaving 529,018 SNPs for GWAS. SNPs with a Bonferroni-adjusted p-value <0.05 were considered significant. Due to missing phenotypic data, not all 942 individuals were included in mapping. For manual traits other than TW, 791 unique individuals were included in GWAS. For image-based traits and TW, 660 individuals were included in GWAS.

The PHW65 NAM was mapped using stepwise model selection with the GBS SNPs followed by resampling GWAS with the 10.6 million projected parental SNPs, as described in Tian *et al*. 2011. The stepwise model selection step, performed using the StepwiseAdditiveModelFitterPlugin in TASSEL5 (Bradbury *et al*. 2007), added SNPs below an entry threshold (set by permutation for each trait, ranging from 9.6×10^-5^ to 1.5×10^-4^) and allowed terms to exit the model based on a limit set at twice the entry threshold. Residuals from the final stepwise model were calculated separately for each chromosome, with any SNPs on that chromosome excluded from the model. The residuals for each chromosome were used as the dependent variable in a resampling-based mapping method similar to that described in Valdar *et al*. 2006 and Tian *et al*. 2011, implemented with the ResamplingGWASPlugin in TASSEL5 (Bradbury *et al*. 2007). Briefly, 80% of the PHW65 NAM individuals were randomly selected and forward regression was used to fit SNPs that were associated with the residuals at a significance level less than 1×10^-4^. This process was repeated 100 times for each combination of trait and chromosome, and each SNP fitted by the resampling GWAS method was assigned a “resampling model inclusion probability” (RMIP) between 0 and 1 reflecting the proportion of models in which that SNP was included. A permutation-based threshold for RMIP was calculated in a manner similar to that described in Tian *et al*. 2011 and five permutations were performed for each chromosome, for each trait. An RMIP threshold of ≥0.05 kept the number of false positives to ≤5 per trait, or on average, ≤1 false positive genome-wide per permutation.

### Selection statistics

We used the 41 BSSSC0 and 21 ex-PVP lines to perform genome-wide scans for selection. The BSSSC0 lines represent maize stiff stalk germplasm from the early 20^th^ century, while the ex-PVP lines represent the product of approximately 60 years of commercial maize breeding. We calculated XP-EHH (Sabeti *et al*. 2007) between the two groups using the WiDiv-942 SNPs with hapbin (Maclean *et al*. 2015), with the –minmaf argument set to 0 and the –scale argument set to 20,000bp. Only SNPs with no missing data in either subpopulation were used. Scores were binned by taking most extreme XP-EHH score from each non-overlapping 10kb window.

XP-CLR (Chen *et al*. 2010) was calculated with XPCLR v1.0 (https://reich.hms.harvard.edu/sites/reich.hms.harvard.edu/files/inline-files/XPCLR.tar) with a grid size of 100bp, a genetic window size of 0.1cM, a maximum of 50 SNPs per window, and correlation level of 0.7. The resulting values were also binned by using the maximum score in each non-overlapping 10kb window to represent that window.

F_ST_ between the two groups was calculated using http://beissingerlab.github.io/docs/vectorFst.R (Beissinger *et al*. 2014), with a correction for small number of populations and uneven sample sizes according to Weir and Cockerham (1984). As with XP-EHH and XP-CLR scores, F_ST_ values were binned into 10kb windows represented by the maximum score in that window.

XP-EHH, XP-CLR, and F_ST_ scores were assigned to each SNP significantly associated with each of the 15 traits by using the score for the 10kb window to which each SNP belonged. Because the distributions of selection statistics for tassel-associated SNPs could be affected by having certain SNPs physically near each other or represented more than once (associated with more than one trait), we used 10,000 iterations of a circular permutation method (Wallace *et al*. 2014) to create a null distribution for each selection statistic. Circular permutation is done by keeping the order and relative position of all the hits intact, while randomizing their starting position along the chromosome. The relationships between the tassel-associated SNPs are maintained, but their relationship to selection statistics along the chromosome is randomized.

The distributions of selection statistics for the tassel-associated SNPs were compared to the circular permutation null distributions both visually and using the Kolmogorov-Smirnov test with a two-sided p-value (Колмогоров 1933; Smirnov 1948).

We also investigated whether the distribution of selection statistics for tassel-associated SNPs was being driven by traits characteristic of the ex-PVPs or the BSSSC0 inbreds. TL, TLp, SL, SLp, BZ, and SP are all related to the length of the tassel and, with the exception of BZ, have higher values in the ex-PVPs. BN, BNp, BD, CP, FD, SK, PR, TW, and TWp are more related to the quantity and density of branches, and have higher values in the BSSSC0 lines. The distributions of XP-EHH scores for these “length” and “branchiness” traits were examined separately to see if one or the other type of trait had undue influence on the overall distribution.

### Data Availability

Tassel images and genotypic data for the PHW65 NAM are available on the CyVerse Data Store (*DOI requested; still pending*) and genotypic data for the WiDiv-942 can be found at doi: 10.5061/dryad.n0m260p (still pending activation; Mazaheri et al., in press). Code is available on GitHub at github.com/joegage/tassel_selection. Supplemental files are available at Figshare. File S1 contains inbred line names and BSSSC0, Public, or ex-PVP designations used for phenotypic comparisons and selection scans. File S2 contains positions of all GWAS hits, along with the 10kb window in which they are located and the selection statistic scores for those windows. File S3 contains BLUP values of WiDiv-942 and PHW65 NAM individuals for 16 tassel morphological triats.

## Acknowledgments

The authors would like to thank Dustin Eilert and Marina Borsecnik for their immense contribution in organizing and conducting field trials at the University of Wisconsin, Madison, as well as Scott Stelpflug, Jonathan Renk, Brett Burdo, Katie Csizmadia, and Mike Ziehr for their help with tassel image acquisition. Image analysis was performed using the compute resources and assistance of the UW-Madison Center for High Throughput Computing (CHTC) in the Department of Computer Sciences. The CHTC is supported by UW-Madison, the Advanced Computing Initiative, the Wisconsin Alumni Research Foundation, the Wisconsin Institutes for Discovery, and the National Science Foundation, and is an active member of the Open Science Grid, which is supported by the National Science Foundation and the U.S. Department of Energy’s Office of Science.

## Author Contributions

Data analysis: J.L.G and M.R.W; Materials development and curation: J.W.E, S.M.K, N.d.L; Manuscript preparation: J.L.G, M.R.W, J.W.E., S.M.K., N.d.L.

## Supplemental Figures

**Supplemental Figure 1:**
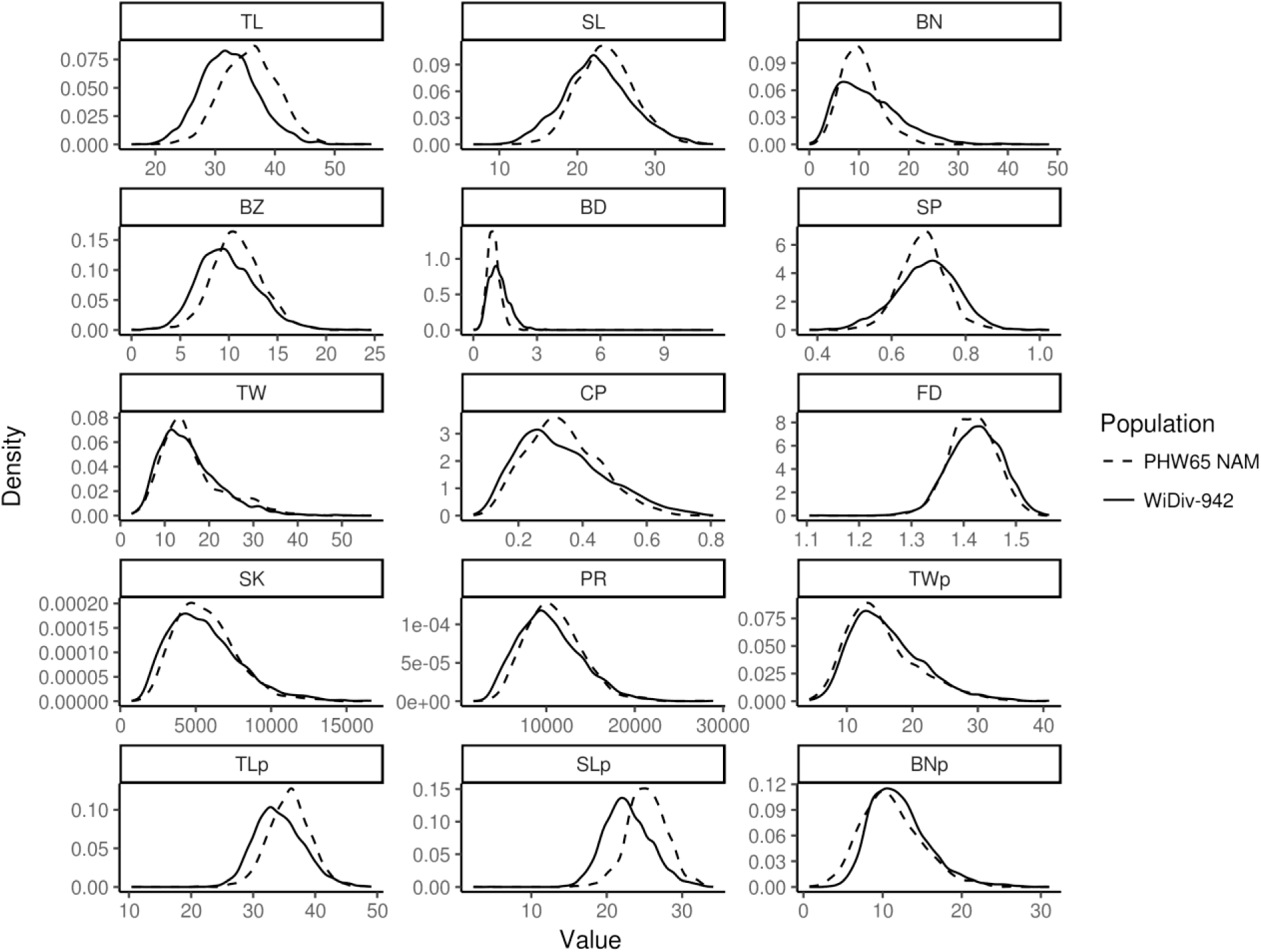
Trait distributions. Distributions of all raw phenotypic values for the WiDiv-942 (solid) and PHW65 NAM (dashed).

**Supplemental Figure 2:**
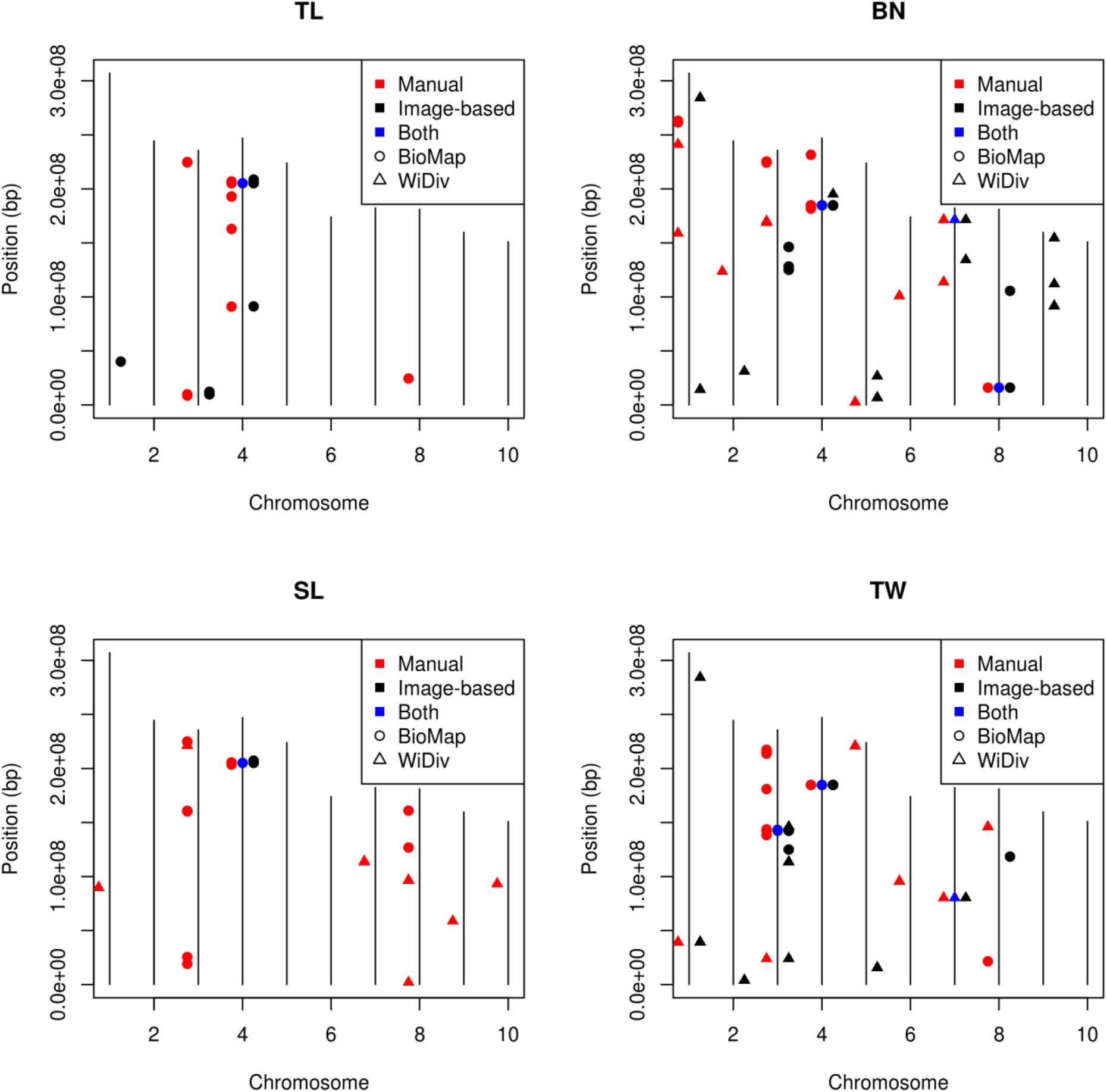
Colocalization of manual and image-based GWAS hits. Tassel length (TL), spike length (SL), branch number (BN), and tassel weight (TW) were all measured manually and by image-based methods. Red and black points show locations of GWAS hits from manual and image-based traits, respectively. If both measurements identified the same exact SNP, the SNP is additionally marked by a blue point. Horizontal ticks represent known inflorescence development genes.

**Supplemental Figure 3:**
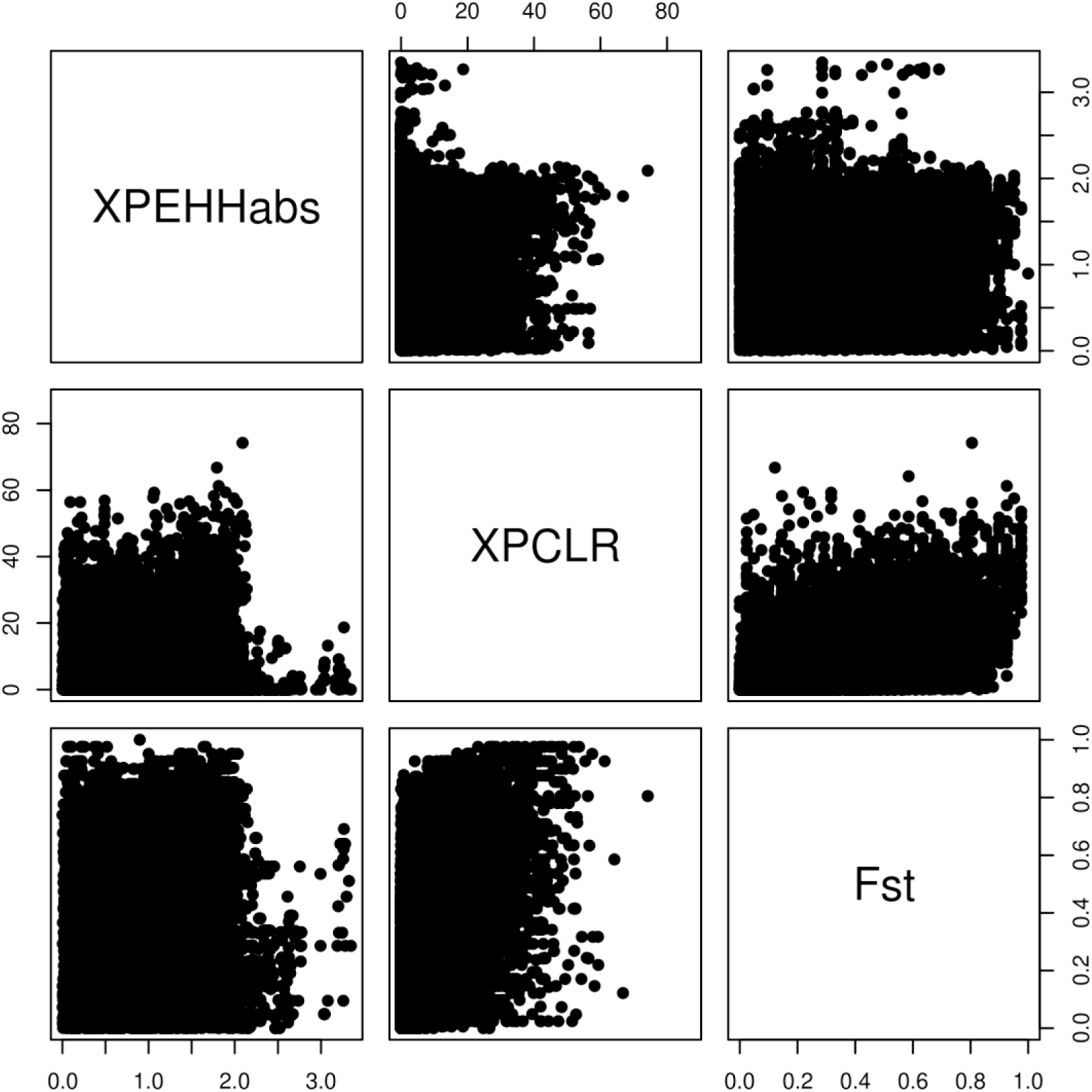
Genome-wide relationships between selection statistics. Comparison of XP-EHH, XPCLR, and F XP-EHH, XPCLR, and F_ST_ values for all windows shows little systematic relationship between statistics. XPEHHabs represents the absolute value of XP-EHH.

**Supplemental Figure 4:**
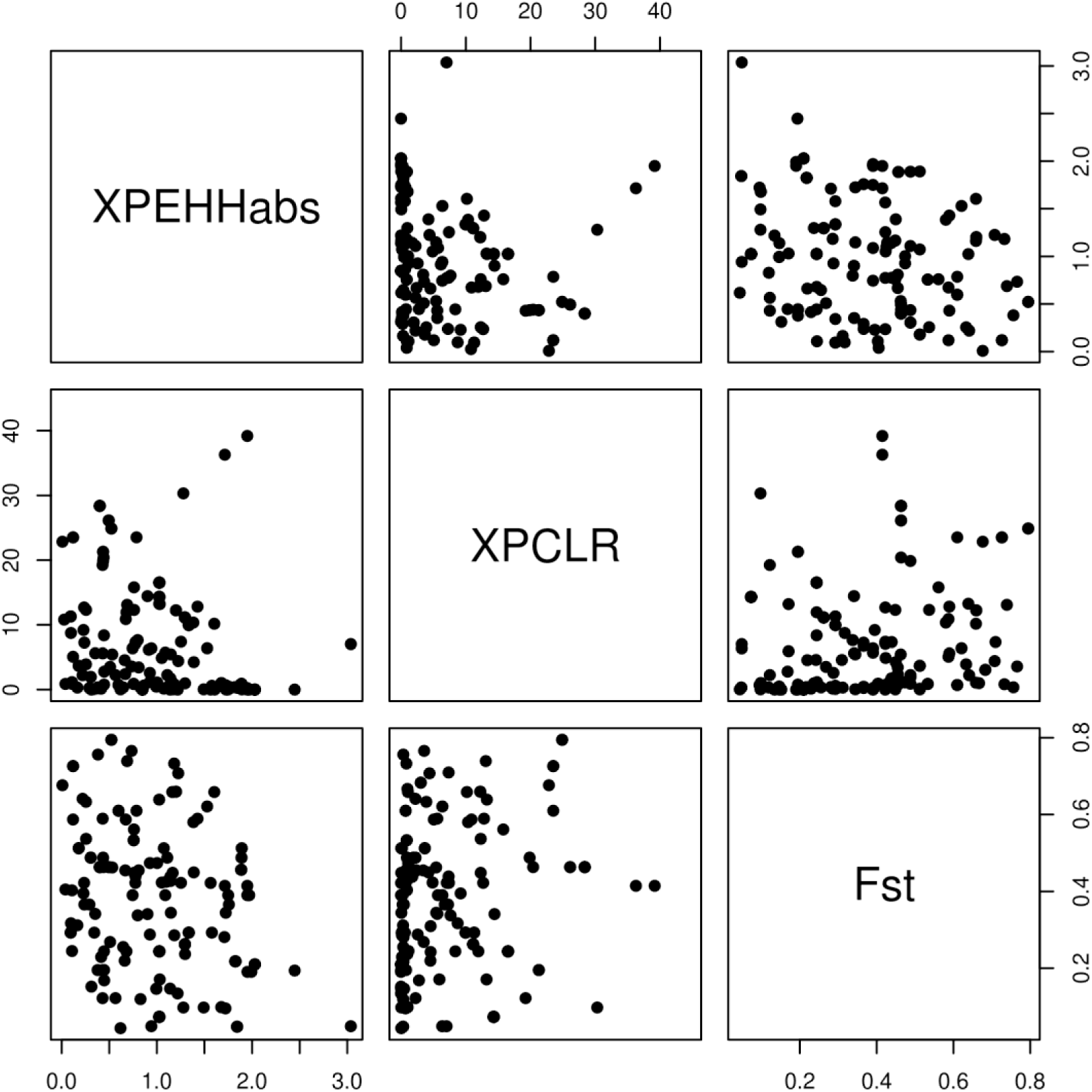
Relationships between selection statistics at GWAS hits. Comparison of XP-EHH, XPCLR, and F XP-EHH, XPCLR, and F_ST_ values for all windows containing SNPs significantly associated with morphological traits shows little systematic relationship between statistics. XPEHHabs represents the absolute value of XP-EHH.

